# CRISPR-Cas12a/Cpf1-assisted precise, efficient and multiplexed genome-editing in *Yarrowia lipolytica*

**DOI:** 10.1101/847905

**Authors:** Zhiliang Yang, Harley Edwards, Peng Xu

## Abstract

CRISPR-Cas9 has been widely adopted as the basic toolkit for precise genome-editing and engineering in various organisms. Alternative to Cas9, Cas12 or Cpf1 uses a simple crRNA as a guide and expands the protospacer adjacent motif (PAM) sequence to TTTN. This unique PAM sequence of Cpf1 may significantly increase the on-target editing efficiency due to lower chance of Cpf1 misreading the PAMs on a high GC genome. To demonstrate the utility of CRISPR-Cpf1, we have optimized the CRISPR-Cpf1 system and achieved high-editing efficiency for two counter-selectable markers in the industrially-relevant oleaginous yeast *Yarrowia lipolytica*: arginine permease (93% for CAN1) and orotidine 5’-phosphate decarboxylase (∼96% for URA3). Both mutations were validated by indel mutation sequencing. For the first time, we further expanded this toolkit to edit three sulfur house-keeping genetic markers (40%–75% for *MET2*, *MET6* and *MET25*), which confers yeast distinct colony color changes due to the formation of PbS (lead sulfide) precipitates. Different from Cas9, we demonstrated that the crRNA transcribed from a standard type II RNA promoter was sufficient to guide Cpf1 endonuclease activity. Furthermore, modification of the crRNA with 3’ polyUs facilitates the faster maturation and folding of crRNA and improve the genome editing efficiency. We also achieved multiplexed genome editing, and the editing efficiency reached 75%–83% for duplex genomic targets (*CAN1-URA3* and *CAN1-MET25*) and 41.7% for triplex genomic targets (*CAN1-URA3-MET25*). Taken together, this work expands the genome-editing toolbox for oleaginous yeast species and may accelerate our ability to engineer oleaginous yeast for both biotechnological and biomedical applications.

## Introduction

*Y. lipolytica*, as a promising oleaginous yeast cell factory, has been extensively engineered for the production of lipids (Qiao, Wasylenko et al. 2017, Xu, Qiao et al. 2017), oleochemicals (Xu, Qiao et al. 2016), drop-in transportation fuels (Xu, Qiao et al. 2016) and commodity chemicals (Blazeck, Hill et al. 2015, Wang, Zong et al. 2019) recently. It is known as a ‘generally regarded as safe’ (GRAS) organism for the production of organic acids and polyunsaturated fatty acids (PUFAs) (Bailey, Madden et al. 2010, Sharpe, Rick et al. 2014) in the food and nutraceutical industry. Compared to *S. cerevisiae, Y. lipolytica* lacks Crabtree effects, without generation of ethanol under high glucose conditions. The low pH tolerance (Cui, Gao et al. 2017), strictly aerobic nature (Abghari and Chen 2014, Ledesma-Amaro, Lazar et al. 2016) and versatile substrate-degradation profile (Ledesma-Amaro, Lazar et al. 2016, Li and Alper 2016, Rodriguez, Hussain et al. 2016) enable its robust growth from a wide range of renewable feedstocks, including pentose (Ledesma-Amaro, Lazar et al. 2016, Li and Alper 2016, Rodriguez, Hussain et al. 2016), crude glycerol (Gao, Yang et al. 2016, Dimitris, Zoe et al. 2019), glacial acetic acids (Xu, Liu et al. 2017, Liu, Marsafari et al. 2019) and volatile fatty acids (VFA) (Spagnuolo, Shabbir Hussain et al. 2018) *et al*. Unlike bacteria, the spatially-organized subcellular compartment and hydrophobic lipid bodies in oleaginous yeast provide the ideal environment for the regioselectivity and stereoselectivity of many plant-specific P450 enzymes (Lv, Koffas et al. 2019). Due to the strong endogenous acetyl-CoA and malonyl-CoA flux, *Y. lipolytica* has been harnessed as an industrial workhorse for efficient synthesis of complex plant secondary metabolites including polyketides (Markham, Palmer et al. 2018, Liu, Marsafari et al. 2019), flavonoids (Lv, Koffas et al. 2019), carotenoids (Gao, Tong et al. 2017, Larroude, Celinska et al. 2018) and terpenoids (Jin, Zhang et al. 2019).

Phylogenetically distant from Baker’s yeast and *S. Pombe, Y. lipolytica* carries 6 chromosomes with 57%∼59% GC content in the coding sequence and a total genome size of 20.5 Mb (Barth and Gaillardin 1997). Compared to *Rodosporidium toruloides*, the low intron density (0.17) in *Y. lipolytica* (Mekouar, Blanc-Lenfle et al. 2010) allows us to easily modify its endogenous pathway and repurpose lipogenesis for various applications. Genome annotation indicates this yeast contains more than 200 hydrophobic compounds assimilation pathways associated with alkane uptake, lipid oxidation and VFA detoxification (Fickers, Benetti et al. 2005) *et al*. This feature makes this yeast a superior host to utilize recalcitrant waste/toxic substrates for eco-friendly production of green chemicals (Liu, Marsafari et al. 2019). Due to its prominent industrial potential, a significant amount of work has been focused to develop genetic toolbox in this yeast, ranging from protein expression (Juretzek, Le Dall et al. 2001, Nicaud, Madzak et al. 2002, Bordes, Fudalej et al. 2007), promoter characterization (Blazeck, Liu et al. 2011, Blazeck, Reed et al. 2013, Liu, Marsafari et al. 2019), gene deletions (Bredeweg, Pomraning et al. 2017, Jang, Yu et al. 2018), YaliBrick-based cloning (Wong, Engel et al. 2017, Wong, Holdridge et al. 2019), Golden-gate cloning (Celinska, Ledesma-Amaro et al. 2017, Larroude, Celinska et al. 2018), Piggybac transposon (Wagner, Williams et al. 2018), iterative gene integration (Gao, Tong et al. 2017, Lv, Edwards et al. 2019) to CRISPR/Cas9-mediated genome-editing (Schwartz, Hussain et al. 2016, Wong, Engel et al. 2017, Holkenbrink, Dam et al. 2018, Morse, Wagner et al. 2018) *et al*. This genetic toolbox affords us a collection of facile genetic tools for streamlined and accelerated pathway engineering in oleaginous yeast species.

Despite that the *Streptococcus pyogenes* (Sp) CRISPR-Cas9 system has been widely adapted for genome editing in multiple yeast species, including bakers’ yeast (DiCarlo, Norville et al. 2013, Jakočiūnas, Bonde et al. 2015, Mans, van Rossum et al. 2015, Si, Chao et al. 2017), the pathogenic yeast *Candida albicans* (Vyas, Barrasa et al. 2015, Min, Ichikawa et al. 2016, Ng and Dean 2017, Shapiro, Chavez et al. 2018), the basidiomycetous yeast *Rodosporidium toruloides* (Jiao, Zhang et al. 2019, Otoupal, Ito et al. 2019, Schultz, Cao et al. 2019), and oleaginous yeast *Y. lipolytica* (Schwartz, Hussain et al. 2016, Wong, Engel et al. 2017, Holkenbrink, Dam et al. 2018, Morse, Wagner et al. 2018) *et al*, the adoption of the recently discovered CRISPR-Cas12/cpf1 (Zetsche, Gootenberg et al. 2015, Zetsche, Heidenreich et al. 2017, Chen, Ma et al. 2018) as genome-editing tools in oleaginous yeast has been significantly lagging behind. The CRISPR-Cpf1 system is complementary to Cas9 with several advantages. Cas9 predominantly recognizes purine rich PAMs (NGG) and cleaves target DNA upstream of PAMs to generate blunt-end DSBs (double strand breaks) (Cong, Ran et al. 2013). In contrast, Cpf1 primarily recognizes T-rich PAMs (TTTN) and cuts DNA in a staggered pattern downstream of the targeted sequence to generate sticky ends (Fig. 1), leaving behind 4∼5 nt 5’-overhangs (Zetsche, Gootenberg et al. 2015). NHEJ (non-homologous end joining repair) usually destroys the PAM site in Cas9 cutting due to its close proximity to the cleavage site, thus preventing future edits and making it difficult for Cas9 re-targeting. Unlike Cas9, Cpf1 cleaves the target DNA 18∼23 nt downstream of the PAM. During NHEJ repair with Cpf1 cutting, the PAM site will be retained and Cpf1 may perform repeated editing (or second chance editing) for the same genetic target. The repeated cutting of Cpf1 may enrich the cleavage events and improve its on-target editing efficiency. Efficient cutting in Cas9/sgRNA also requires the proper folding of tracrRNA (trans-activating CRISPR RNA) and crRNA. Unlike Cas9, Cpf1 only use a 20 nt DR (directed repeat) sequence preceding the crRNA. Without the 80 nt tracrRNA, this simple crRNA in Cpf1 allows us to deliver the guide RNA much more efficiently than we can deliver the sgRNA in the Cas9 system. The expanded PAM sequence space, staggered cutting patterns and the simplified crRNA structures make Cpf1 an attractive complementation to enable broader and improved targeting opportunities for oleaginous species.

**Fig. 1.**
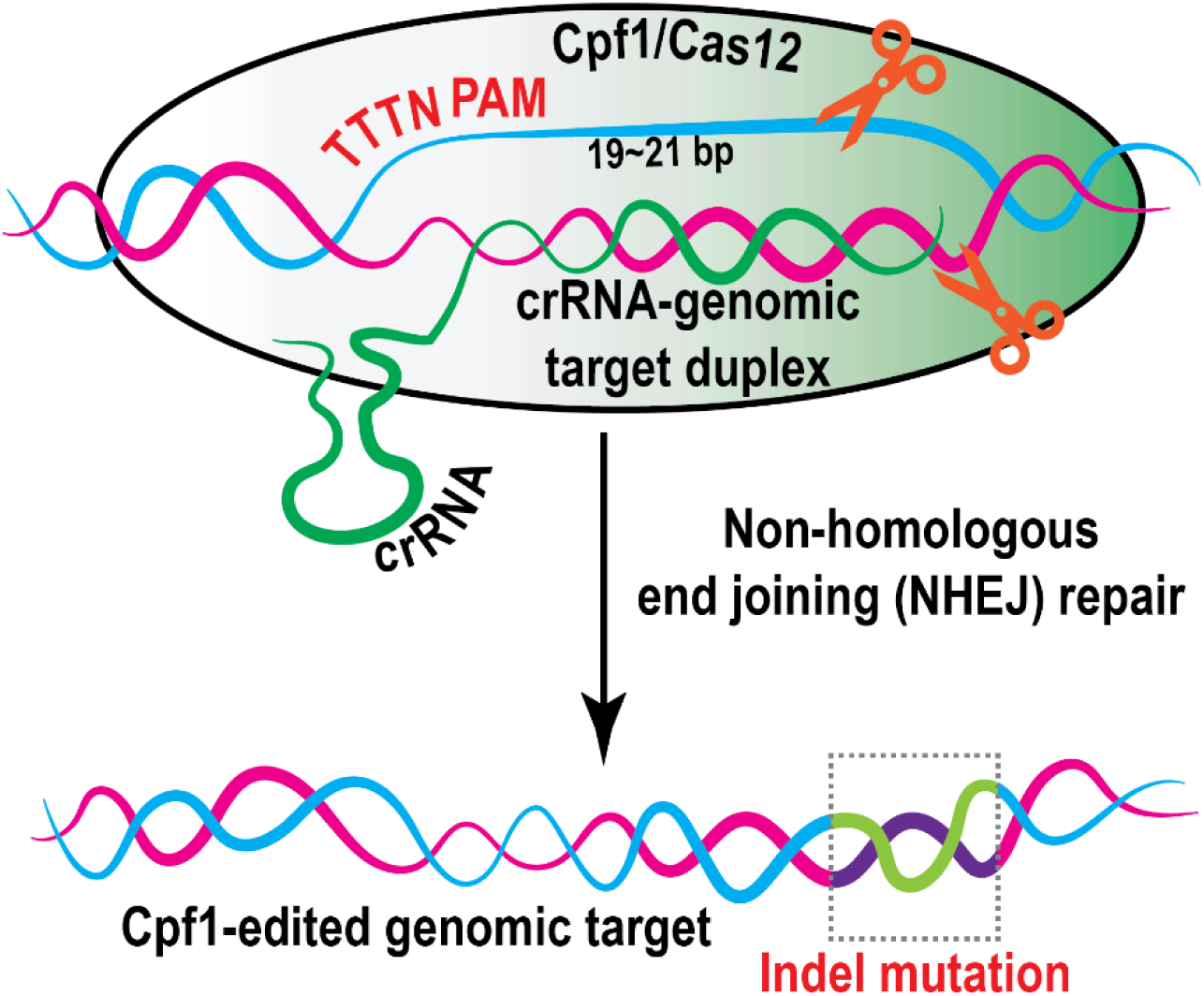
Crispr-cas12/cpf1 recognizes TTTN protospacer adjacent motif (PAM) and is guided by a simple crRNA without the need for tracrRNA (trans-activating CRISPR RNA). Cpf1 introduces double strand break (DSB) in a staggered pattern and generate sticky ends. The DSB is primarily repaired by non-homologous end joining (NHEJ) in *Y. lipolytica*.

In this work, we demonstrated that AsCpf1 could efficiently introduce nucleotide substitutions, insertions and gene deletions with high efficiency and accuracy in a multiplexed manner. Various promoters ranging from type II promoter flanked with ribozymes, type II promoter without ribozymes, type III hybrid SCR1’-tRNA^Gly^ promoter and the 5sRNA promoter were tested to optimize crRNA expression. Our engineered CRISPR-Cpf1 system has been used to edit two counter-selectable genetic markers, URA3 (encoding orotidine 5’-phosphate decarboxylase) and CAN1 (encoding arginine permease). This system was further applied to edit three visual-selectable genetic markers encoding the sulfur-housekeeping genes (MET2, MET6 and MET25) of *Y. lipolytica*, with high cleavage efficiency. We further performed multiplexed targeting by using a tandem array of three crRNAs for URA3, CAN1 and MET25 and achieved high targeting efficiency for duplex and triplex editing. Taken together, this CRISPR-Cpf1-assisted system provides a highly efficient and versatile toolkit that expanded our capability for genome-targeting and engineering in oleaginous yeast species.

## Results and Discussions

### CRISPR-Cpf1 mediated in-del mutations of counter-selectable marker CAN1

As a proof-of-concept to test the genome-editing efficiency of the CRISPR-Cpf1 system in *Y. lipolytica*, we first targeted the *CAN1* gene (encoding arginine permease). Arginine permease (CAN1) is responsible for the yeast to assimilate arginine from the media. Anti-metabolite, L-canavanine is a structural analog to arginine and will stop polypeptide synthesis if the cell mistakenly takes up canavanine. *CAN1* mutation confers resistance to L-canavanine toxicity, which allows counter-selection of *CAN1* mutants on CSM-Arg agar plates supplemented with L-canavanine. In order to implement the CRISPR-Cpf1 system in *Y. lipolytica*, the AsCpf1 gene was codon-optimized using *homo sapiens* codon usage (Zetsche, Gootenberg et al. 2015) and expressed using the strong TEF-intron promoter (Fig. 2A). A nuclear localization signal (NLS) was fused to the C-terminal of AsCpf1 to localize the Cas12/Cpf1 endonuclease to the nucleus. The TEF-intron promoter has been characterized as a strong constitutive promoter and employed to express the sgRNA to implement CRISPR-Cas9 based editing of *CAN1* in *Y. lipolytica* (Wong, Engel et al. 2017). We first expressed the crRNA of *CAN1* using this well-characterized TEF-intron promoter. The crRNA sequence was flanked by the hammer head ribozyme (HHR) and hepatitis delta virus (HDV) ribozyme (Bayer and Smolke 2005, Gao and Zhao 2014, Gao, Tong et al. 2016) to facilitate the release of functional crRNA from primary transcript via self-cleavage of the two ribozymes (Fig. 2B). The CRISPR-Cpf1 mediated DSB was primarily repaired by non-homologous end joining (NHEJ) in *Y. lipolytica*. Colony PCR products were sequenced to identify in-del mutations at the target site. As was shown in Fig. 2D, random insertion or deletion mutations were detected at the crRNA target site downstream of the TTTN PAM in all the canavanine-resistant colonies. Most strikingly, the editing efficiency of *CAN1* in *Y. lipolytica* po1g strain reached 72.9%±9.5% (12/16, 13/16, 10/16) using this crRNA expression strategy (Fig. 2C), which achieved higher editing-efficiency than that of using Cas9 with the same crRNA expression strategy in our previous work (Wong, Engel et al. 2017, Wong, Holdridge et al. 2019).

**Fig. 2.**
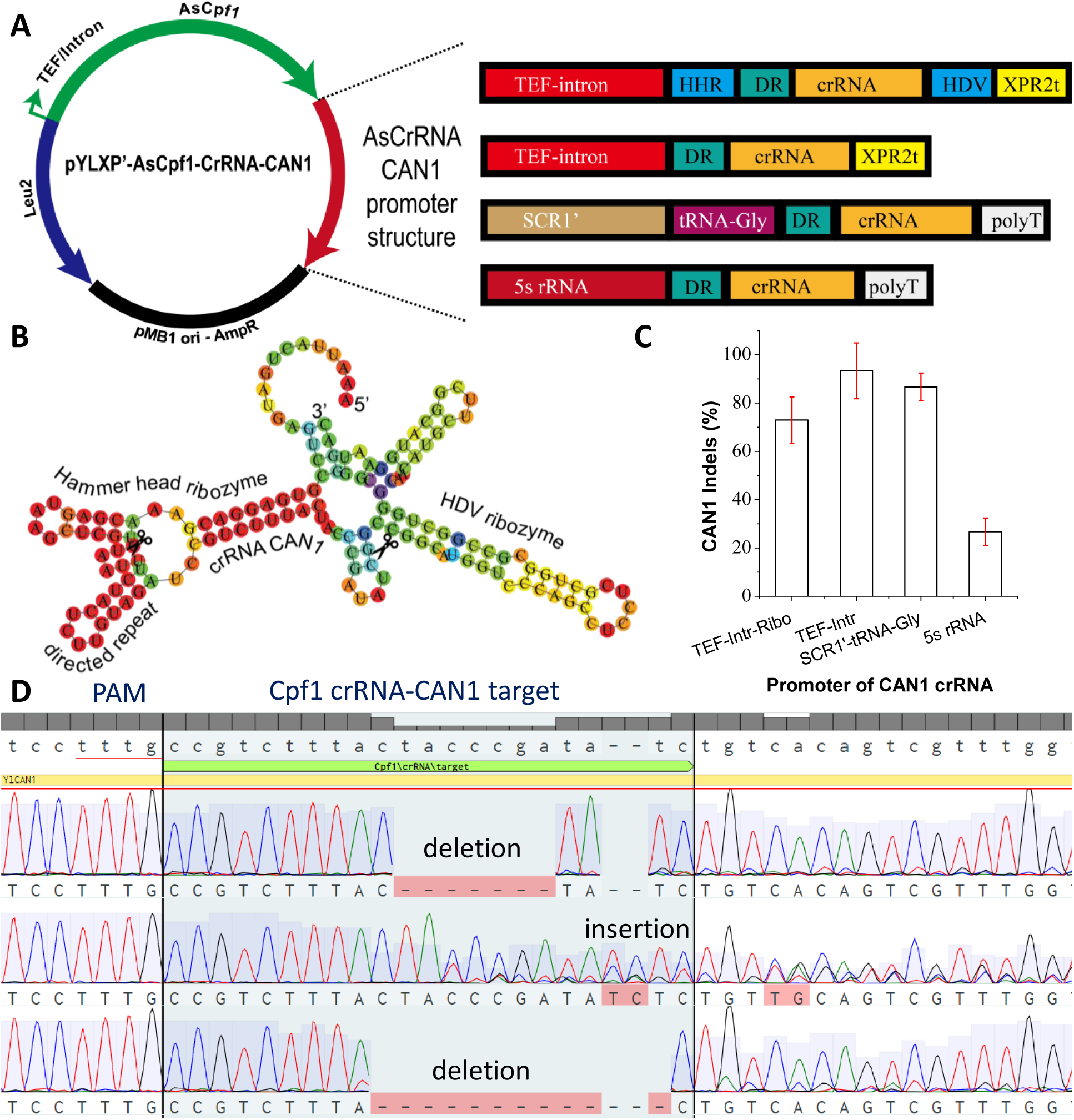
Crispr-cas12/cpf1-mediated genome-editing of dominant marker arginine permease encoded by *CAN1*. (**A**) Four genetic configurations of crRNA promoter were tested to drive the expression of crRNA-CAN1. (**B**) Type II RNA promoter (TEF) were flanked with upstream hammer head ribozyme and downstream HDV ribozyme to drive the expression of crRNA-CAN1. Ribozyme self-cleavage site has been marked with a scissor. (**C**) On-target genome-editing efficiency for CAN1 with four crRNA promoter configurations. *N=3, the reported data represents mean* ± *sd*. (**D**) Sanger DNA sequencing to validate the on-target insertion or deletion (indels) mutations for *CAN1* loci.

Genome editing of *CAN1* gene has been reported in other studies. Bao *et al*. described the editing of *CAN1* ranging from 14.71% using a 100 bp donor DNA to 100% when prolonged incubation in liquid media was performed in *S. cerevisiae* (Bao, Xiao et al. 2014). Similarly, in-del mutation of *CAN1* in *Y. lipolytica* was only found to be 7∼10% in the CRISPR-Cas9 system (Wong, Engel et al. 2017, Wong, Holdridge et al. 2019), when the sgRNA was flanked with 5’ HHR and 3’ HDV from the strong TEF/intron promoter. In a recent study, targeting efficiency of *CAN1* by Cas9 achieved 60% using a T7 promoter in *Y. lipolytica* (Morse, Wagner et al. 2018). Compared with extant strategies, the CRISPR-AsCpf1 system demonstrated superior editing efficiency to introduce in-del mutations in *Y. lipolytica*, and prolonged incubation was not necessary. Potentially, this CRISPR-Cpf1 system could be implemented in a relatively shorter timescale with high editing efficiency.

### Optimizing crRNA expression to improve CRISPR-Cpf1 targeting efficiency

CRISPR RNA (crRNA) belongs to non-coding RNA and transcription of crRNA was found to be critical for CRISPR genome editing. A modified type II promoter such as the TEF promoter or a type III promoter such as the hybrid SNR52 promoter or the U6 promoter, is generally used to drive the expression of sgRNA or crRNA. We demonstrated that the type II promoter (TEF-intron with ribozymes) could achieve relatively high genome-editing efficiency (Fig. 2). To provide the optimal crRNA transcript, we constructed three additional crRNA constructs for *CAN1* targeting (Fig. 2A): (I) TEF-intron promoter *<u>without</u>* 5’HHR or 3’ HDV ribozymes; (II) the hybrid promoter SCR1’-tRNA^Gly^ (Schwartz, Hussain et al. 2016) and (III) the 5s rRNA promoter (Schultz, Cao et al. 2019). Interestingly, the crRNA expressed by TEF-intron promoter without ribozymes reached 93.3%±11.5% (10/10, 10/10, 8/10) editing efficiency (Fig. 2C). This indicated that Cpf1 alone is sufficient to process crRNA and generate mature crRNA for Cpf1 targeting, which is consistent with recent findings that Cpf1 was able to process pre-crRNA arrays to form mature crRNA (Fonfara, Richter et al. 2016, Zetsche, Heidenreich et al. 2017). In *Pichia pastoris*, polymerase II promoter PHXT was also recently used for the expression of sgRNA in CRISPR-Cas9 based system recently (Weninger, Hatzl et al. 2016). It was found that direct fusion of the sgRNA to PHXT led to high targeting efficiency.

An editing efficiency of 86.6%±5.7% (9/10, 9/10, 8/10) was also achieved for *CAN1* targeting using the hybrid SCR1’-tRNA^Gly^ promoter (Fig. 2C), indicating that type III RNA polymerase supports the expression and processing of functional crRNAs. In a recent study, the short and abundant 5s ribosomal RNA (rRNA) was employed to drive the expression of sgRNA in *Aspergillus niger* with high genome-editing efficiency (Zheng, Zheng et al. 2018). To expand this work, we also tested the feasibility of using the native promoter from *Y. lipolytica*, 5s rRNA, to express the cRNA of *CAN1*. The editing efficiency of *CAN1* under the 5s rRNA promoter only reaches 10%-20% (Fig. 2C), indicating the complex non-coding RNA processing mechanism across different microbial hosts.

In summary, the type II TEF-intron promoter (without ribozymes) and the hybrid SCR1’-tRNA^Gly^ promoter were identified as the most efficient promoters to drive the expression of crRNA in this CRISPR-Cpf1 system, leading to consistently high in-del editing of *CAN1* up to 80%∼90%. This efficiency represents a significantly improved gene-editing compared to previously reported CRISPR-Cas9 systems in oleaginous yeast species (Gao, Tong et al. 2016, Wong, Engel et al. 2017, Wong, Holdridge et al. 2019).

### CRISPR-Cpf1 mediated in-del mutations of counter-selectable marker URA3

We further tested another counter-selectable genetic marker to validate the functionality of CRISPR-Cpf1 in *Y. lipolytica*. The *URA3* gene (encoding orotidine 5’-phosphate decarboxylase) is a commonly-used yeast genetic marker. *URA3* mutants will require the supplementation of uracil, but develop resistance to the toxic 5’-fluoroorotic acid (5’-FOA). Since our host strain *Y. lipolytica* po1f carries a non-functional allele of *URA3* gene (the endogenous *URA3* was partially disrupted). To minimize the mis-targeting on the endogenous URA3 remnants, we first modified the po1f strain by replacing the *ALK7* (encoding alkane oxidase) gene with a well-defined *URA3* gene (supplementary Fig. 1). This strain was subsequently used as the host for genome editing of *URA3*. Two types of crRNAs, URA3A and URA3B, targeting either the sense or the anti-sense strand, respectively, were designed and expressed using the TEF-intron promoter with ribozymes (Fig. 3A). 5’-FOA resistant colonies were harvested and used as template to amplify the edited *URA3* gene. After sequencing of these PCR fragments, we identified in-del mutations at both the sense strand and nonsense strand (Fig. 3B and Fig. 3C). The hybrid SCR1’-tRNA^Gly^ promoter led to about 100% disruption of *URA3* using crRNA-URA3A. Similar editing efficiency was achieved with the TEF-intron promoter with ribozymes (Fig. 3A). These results reinforce that both polymerase II and III promoters can be used to drive the expression of crRNA in CRISPR-Cpf1 system, albeit editing efficiency may vary depending on the specific genetic targets (For instance, the *CAN1* marker in previous section).

**Fig. 3.**
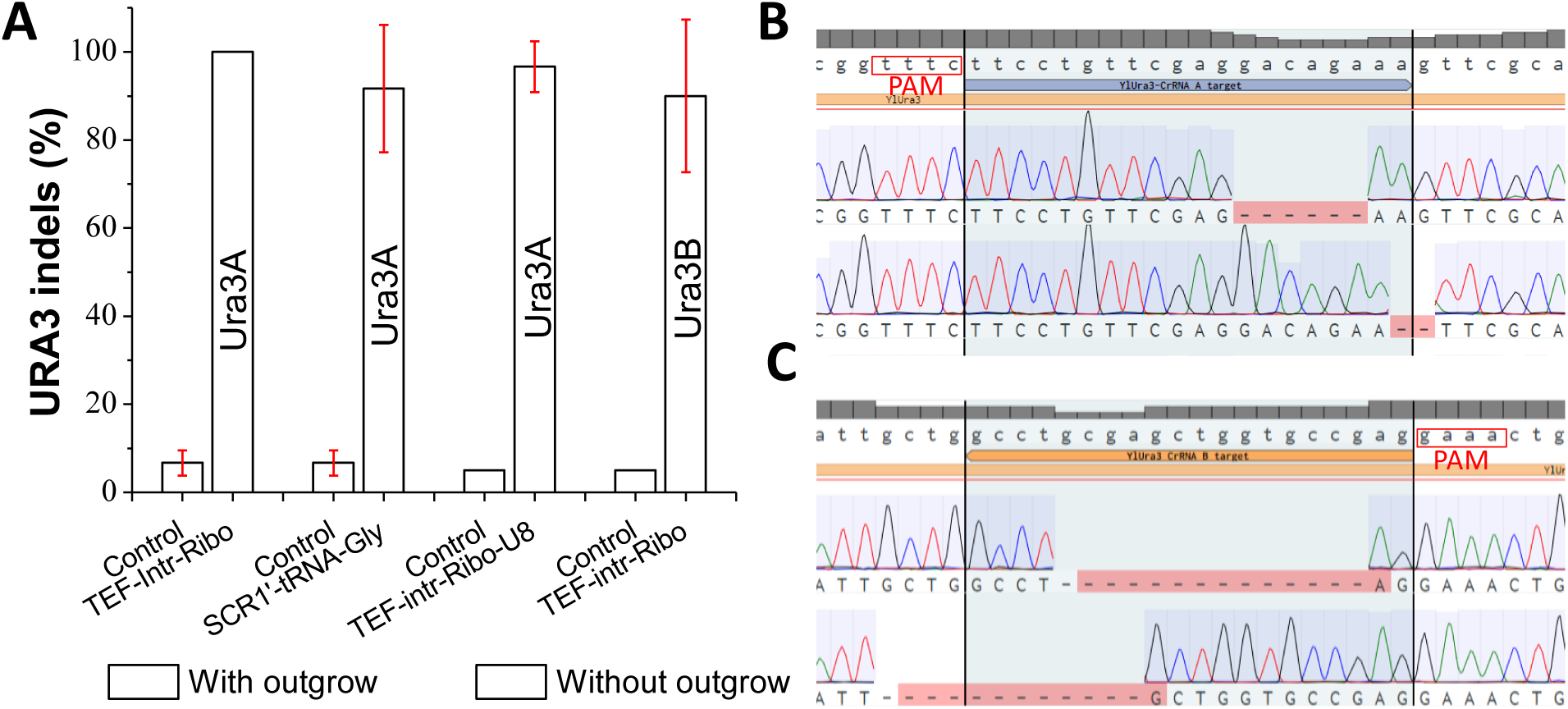
Crispr-cas12/cpf1-mediated genome-editing of counter-selectable marker orotidine 5’-phosphate decarboxylase encoded by *URA3*. (**A**) On-target genome-editing efficiency for URA3 with different crRNA configurations and modifications. *N=3, the reported data represents mean* ± *sd*. (**B**) Indel mutations of URA3 genetic marker when the crRNA-URA3A targets to the sense strand. (**C**) Indel mutations of URA3 genetic marker when the crRNA-URA3B targets to the antisense strand.

For screening and identifying the mutant colonies, we found that outgrowth is critical to enrich the genome-editing events. When transformation mixture was directly plated on CSM-Leu plate supplemented with 50 mg/L uracil and 1 mg/mL 5’-FOA, the CFU (colony forming unit) was drastically reduced. Less than ten colonies were obtained on the plate, possibly due to the high selection pressure of 5’-FOA and the slow cleavage activity of Cpf1: majority of cells were killed by 5’-FOA before they could be edited by Cpf1. It has been recently discovered that the addition of poly-thymidine to the 3’-end of crRNA could improve the AsCpf1 genome editing efficiency (Xie, Minkenberg et al. 2015, Nowak, Lawson et al. 2016, Bin Moon, Lee et al. 2018). We thus modified the crRNA by adding eight thymidine (T8), which will form a poly-U overhang following the 3’-end of the crRNA. CFUs were markedly improved by 10-fold over the original crRNA-URA3A after 2 days of outgrowth in CSM-Leu liquid media. *URA3* editing efficiency reached 96% (10/10, 10/10, 9/10), indicating that the U-rich strategy could enhance the CRISPR-Cpf1 genome editing efficiency in *Y. lipolytica* (Fig. 3A), possibility due to the fast maturation of the crRNA in the presence of polyU. This is not surprising, as the formation of the Cpf1-crRNA-DNA nucleoprotein complex has been recently proposed as the major rate-limiting step for Cpf1 genome-editing (Strohkendl, Saifuddin et al. 2018). Either targeting to the sense strand (Fig. 3B) or the non-sense strand (Fig. 3C), we detected in-del mutations downstream of the TTTCs PAMs. Notably, control experiment was performed by directly plating the transformation mixture onto CSM-Leu plate without outgrowth. We observed that only 5% to 7.5% disruption efficiency was achieved (Fig. 3A) for the control experiment. Outgrowth has been a commonly-used strategy to improve the CRISPR-Cas9 targeting efficiency of genes including TRP1 (Gao, Tong et al. 2016) and PEX10 (Schwartz, Hussain et al. 2016) in *Y. lipolytica*. The URA3 mutants obtained after outgrowth showed different in-del mutations, which indicates that the improved efficiency was not due to enrichment of edited mutants. These results highlighted the critical role of outgrowth to enrich genome editing events for dominant and counter-selectable genetic markers, and the polyU-rich strategy could accelerate the maturation of functional crRNAs.

### CRISPR-Cpf1 mediated in-del mutation of non-dominant markers

The methionine pathway starts with a number of sulfur house-keeping genes that sequentially reduce sulfate (SO_4_^2-^) to sulfite (SO_3_^2-^) and sulfide (S^2-^) (Fig. 4A). Disruption of MET25 has been linked to the accumulation of sulfide (S2-) intracellularly in both *S. cerevisiae* (Cost and Boeke 1996) and *C. albicans* (Viaene, Tiels et al. 2000). The yeast colony will form distinct black precipitate (lead sulfide PbS) if colorless and soluble lead (lead nitrate or lead acetate) salt was supplied to the media. This submissive marker has been used as a basic genetic tool to understand sulfur metabolism and methionine biosynthesis in various hosts.

**Fig. 4.**
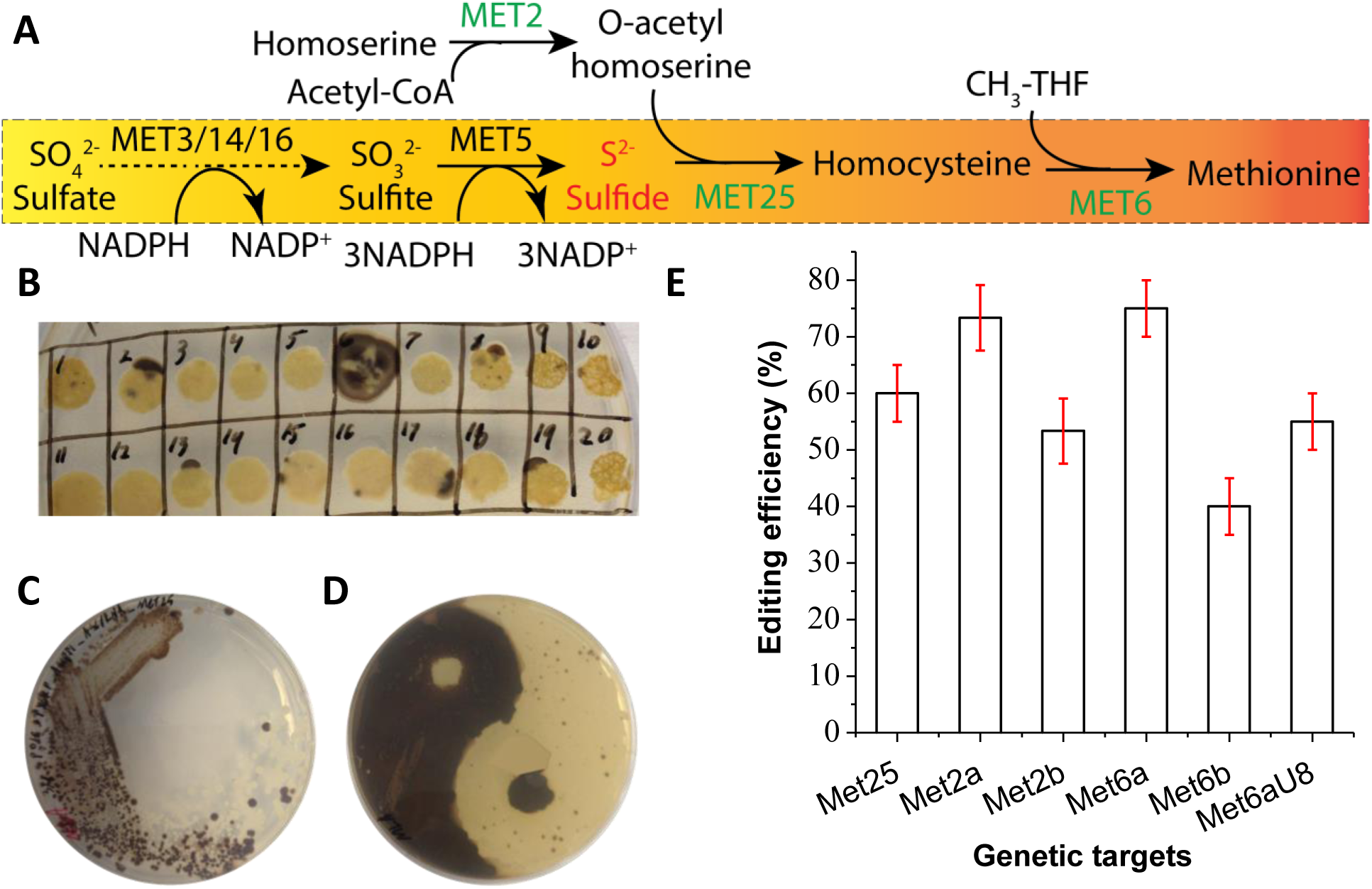
Crispr-cas12/cpf1-mediated genome-editing of submissive marker encoded by sulfur house-keeping genes *MET25, MET6* and *MET2*. (**A**) Methionine biosynthetic pathway is encoded by a number of reduction steps to incorporate sulfide into the carbon backbone of homoserine. Sulfate was sequentially reduced to sulfide, blocking the sulfide incorporation pathway will result in the accumulation of sulfide (S^2-^) intracellularly. (**B**) Agar plate screening of *MET25* indel mutations. Colonies with black sector indicates the successful mutation of *MET25*. (**C**) Re-streaking of the sectored colonies onto MLA plate to isolate genetically pure MET25 mutants. (**D**) A Taichi or “Yin-Yang” art is created by plating the wild type P01g strain and the MET25 mutant strains on the MLA plate. (**E**) On-target genome-editing efficiency for MET25, MET2 and MET6 with different crRNA configurations and modifications. *N=3, the reported data represents mean* ± *sd*.

The genome editing of *CAN1* and *URA3* was achieved by using counter-selectable marker with selection pressure (i.e. canavanine for *CAN1* and 5’-FOA for *URA3*, respectively). To test the editing efficiency of CRISPR-Cpf1 without selection pressure, three submissive markers: *MET25* (encoding homocysteine synthase), *MET2* (encoding O-acetyl homoserine synthase) and *MET6* (encoding homocysteine methyl-transferase) involved in methionine metabolism (Thomas and Surdin-Kerjan 1997) were targeted. After transforming *Y. lipolytica* po1g with the plasmid pYLXP’-AsCpf1-crRNA-MET25, we observed that inactivation of *MET25* indeed led the formation of black or brown colonies (Fig. 4B) in *Y. lipolytica*. Genome editing efficiency of 65% (13/20) was achieved after 7 days of incubation (Fig. 4B). It was also found that the black mutant cells were scattered around the white colonies (Fig. 4C), indicating the genome-editing events are not ubiquitous, and screening is essential to identify the genome-edited cells. To demonstrate the distinct phenotype, a Taichi or Ying-Yang bio-art has been created with the MET25 genome-edited cells and the wild type cells (Fig. 4D).

To test if the disruption of *MET2* and *MET6* could also lead to the black colony, three MET6 crRNA targets (crRNA-MET6A, crRNA-MET6B and crRNA-MET6A-U8) and two MET2 crRNA targets (crRNA-MET2A and crRNA-MET2B) were also designed and assembled into pYLXP’-AsCpf1 plasmid. The precipitation of lead sulfide (PbS) and the formation of black colony were found for all these genetic targets, with varying genome-editing efficiency (Fig. 4E). The disruption efficiency for the three MET6 targets was found to be 75%±5% (16/20, 14/20, 15/20) for Met6a, 40%±5% (8/20, 7/20. 9/20) for Met6b and 55%±5% (10/20, 11/20, 12/20) for Met6aU8, respectively (Fig. 4E). The genome-editing efficiency for the two MET2 targets was found to be 73.3%±5.7% (8/10, 7/10, 7/10) for Met2a, and 53.3%±5.7% (5/10, 6/10, 5/10) for Met2b (Fig. 4E), which is comparable to the editing efficiency obtained from the MET6 marker. After the black colonies were formed, we further detected the *met6* indel mutations with a T7 endonuclease kit, which allows us to identify heteroduplex DNAs containing any mismatches between the two strands of the complementary DNA. Colony PCR was performed and the PCR products were subjected to denaturation and re-annealing to form the heteroduplex DNA. After digestion with T7 endonuclease, two shorter bands corresponding to the cleavage site of the edited PCR products was observed (supplementary Fig. S2), indicating that the genome-editing events occurred at the target site of MET6. Notably, the varying targeting efficiency of the sulfur metabolism genes highlighted the effect of crRNA guide sequence on gene-editing efficiency. For practical applications, several guides should be designed and tested to achieve optimal gene disruption.

### Multiplexed genome-editing with CRISPR-Cpf1

Compared to conventional chromosomal engineering method, including homology-based chromosomal integration (gene knock-in) or inactivation (gene knock-out or deletion), CRISPR provides us the opportunity to precisely edit multiple genomic targets, without the need of iterative marker recycling or curation. This multiplexed genome-editing is commonly achieved by placing an array of crRNAs or sgRNAs (Zhang, Wang et al. 2019). To test whether the CRISPR-Cpf1 system could achieve multiplexed genome editing in *Y. lipolytica*, we first investigated the simultaneous disruption of two genes. The crRNA expression cassettes of CAN1 and MET25 were assembled with the vector containing AsCpf1 (Fig. 5A and supplementary Fig. S3). The double mutants were screened based on canavanine resistance and the formation of black colony. It was found that the efficiency of double mutant of *CAN1* and *MET25* reached 83.3%±5.7% (Fig. 5B). Similarly, the screening of double mutants of *CAN1* and *URA3* was performed based on canavanine resistance and 5’-FOA resistance. The double editing efficiency of *CAN1* and *URA3* was found to be 75%±5% (Fig. 5B). It should be noted that the *URA3* genome-editing is enriched with cell outgrow in uracil-rich media before the cells were screened against 5’-FOA or canavanine resistance.

**Fig. 5.**
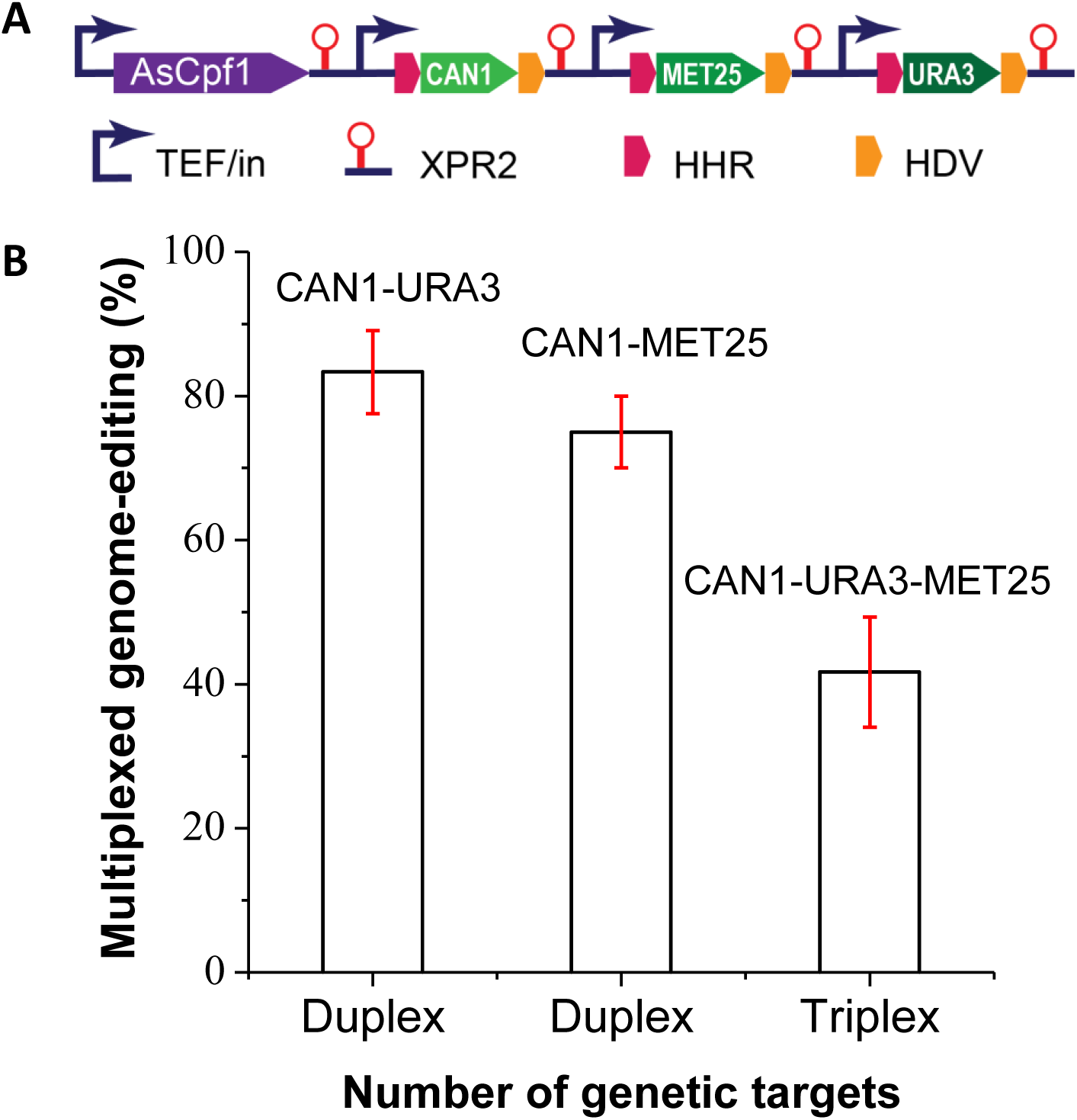
CRISPR-Cpf1/cas12 mediated multiplexed genome-editing for *MET25, URA3* and *CAN1* in *Y. lipolytica*. (**A**) Genetic configurations for three crRNAs arrays to drive multiple genome-editing targets. A detailed plasmid map for this crRNA array could be found in supplementary Fig. S3. (**B**) Duplex genome-editing efficiency for CAN1-URA3 and CAN1-MET25; and triplex genome-editing efficiency for CAN1-URA3-MET25.

We next sought to disrupt three genes with the crRNA targets for *URA3, MET25* and *CAN1*. After cell outgrow in uracil-rich media, the genome-edited cells were screened based on 5’-FOA resistance, the formation of black colony and canavanine resistance (supplementary Fig. S4). We observed that the triplex editing efficiency reached 41.7%±7.6% (Fig. 5B). Despite that this editing efficiency is not high compared to multiplexed genome-editing in *S. cerevisiae* (Zhang, Wang et al. 2019), this efficiency (45%) would guarantee two positive triple mutants out of five colonies, which is sufficient for us to perform genome-editing and strain engineering in this yeast. Nevertheless, further optimization of the crRNA transcripts will be necessary for us to improve the multiplexed genome-editing efficiency.

## Conclusions

In this work, we have tested an alternative genome-editing toolkit, CRISPR-Cas12/Cpf1, to perform genome-editing in *Y. lipolytica*. As a complementary toolbox to CRISPR-Cas9, we have achieved precise, efficient and multiplexed genome-editing with improved cleavage efficiency. The distinct PAM (TTTN) could expand gene-editing at AT-rich regions on genome. In addition, the simplified crRNA structure without the need of tracrRNA makes Cpf1 a potentially better candidate for genome-editing, offering us an easy-to-operate approach to express crRNAs.

With this toolkit, we have tested the CRISPR-Cpf1 genome editing efficiency for two counter-selectable markers (URA3 and CAN1) and three submissive markers (MET2, MET25 and MET6). We have validated genome-editing events (indel mutations) by both DNA sequencing and heteroduplex DNA digestion. To optimize crRNA expression, we also assessed four versions of promoters, ranging from TEF-intron-ribozyme, TEF-intron, tRNA^Gly^ to 5s RNA promoters. Our results demonstrate that TEF-intron promoter alone is sufficient to express crRNAs without ribozymes, possibly due to the RNA-processing ability of AsCpf1. We also observed that modification of crRNAs by adding polyUs downstream of crRNA could facilitate the faster release and maturation of crRNAs from the primary RNA transcripts. For counter-selectable markers (CAN1 or URA3), our CRISPR-Cpf1 editing efficiency was ranging from 72.9% to 96%; for submissive markers (MET2, MET25 and MET6), our editing efficiency was ranging from 40% to 80%. For multiplexed genomic targets, our editing efficiency reached 75∼83% for duplex targets, and 45% for triplex targets. Taken together, this CRISPR-Cpf1 system should complement the Cas9 toolkit and enable us to precisely edit the genomic targets of *Y. lipolytica* with improved editing efficiency. This work provides us an invaluable tool to perform multiplexed genome-editing and may accelerate our ability to deliver oleaginous yeast cell factories for various applications.

## Methods and Materials

### Strains and growth conditions

*E. coli* NEB5α was routinely cultivated in Luria-Bertani (LB) broth and used for all cloning work. *Y. lipolytica* Po1f and Po1g were used as hosts for genome-editing. YPD medium consisting of 10 g/L yeast extract, 20 g/L peptone and 20 g/L Dextrose or complete synthetic media (CSM) lacking proper amino acid were used for yeast cultivation. For the selection of *CAN1* mutants, 50 mg/L L-Canavanine was supplemented in CSM-Arg plates. For the counter-selection of *URA3* mutants, 50 mg/L uracil and 1 mg/mL 5’-fluoroorotic acid (5’-FOA) was added to CSM-Leu agar plates. MLA plates consisting of 3 g/L peptone, 5 g/L yeast extract, 0.2 g/L ammonium sulfate, 40 g/L glucose, 1 g/L lead nitrate and 20 g/L agar were used for visual selection of *MET25, MET2* and *MET6* mutants.

### Plasmid construction

All primers (Supplementary Table S1) used in this work were ordered from Integrated DNA Technologies (USA). All plasmids used in this study were listed in Supplementary Table 2. AsCpf1 was PCR amplified using pY010 as template and AsCpf1_F/AsCpf1_R as primers. The PCR product was designed to retain the nuclear localization signal (NLS) but the 3HA tag was removed. This PCR product was cleaned with Zymoclean kits and Gibson assembled into the SnaB1 and KpnI digested pYLXP’, to yield plasmid pYLXP’-AsCpf1. To construct the crRNA for CAN1 targeting, AvrII-Fwd and Can1-crRR were used to PCR amplify the fragment crRNA-Can1up. SalI_Rvs and Can1-crRF were used to amplify the fragment crRNA-Can1dn. After PCR clean-up, crRNA-Can1up and crRNA-Can1dn were assembled with vector backbone pYLXP’ digested with *Avr*II and *Sal*I using Gibson assembly to yield pYLXP’-AscrRNA-Can1. Other crRNA plasmids were constructed in a similar manner. All plasmids assembled by Gibson assembly were verified by Sanger sequencing (Genewiz, USA). To obtain all-in-one plasmid for genome editing, the crRNA constructs were digested with *Avr*II and *Sal*I. The smaller fragment was gel-recovered and ligated with the *Nhe*I and *Sal*I digested pYLXP’-AsCpf1. All sub-cloned plasmids were verified by double digestion to match the pattern of the software (Benchling) predicted digestion pattern. All strains and plasmids constructed in this work is listed in supplementary Table S2.

### Screening of genome-edited mutants for CAN1,URA3 and MET25/MET2/MET6

Colonies of *Y. lipolytica* Po1g transformed with pYLXP’-AsCpf1-AscrRNA-Can1 were picked up using sterile pipette tips and spotted on CSM-Arg plates supplemented with 50 mg/L L-canavanine. Colonies of *Y. lipolytica* Po1fΔALK7::URA3 transformed with pYLXP’-AsCpf1-AscrRNA-URA3A or pYLXP’-AsCpf1-AscrRNA-URA3B were spotted on CSM-Leu plates with 50 mg/L uracil and 1 mg/mL 5’-FOA. Colonies of *Y. lipolytica* Po1g transformed with pYLXP’-AsCpf1-AscrRNA-Met25 were spotted on MLA plates. For *CAN1* or *URA3* editing, incubation of 2-3 days allowed the formation of colonies on CSM-Arg plates. For *MET25* targeting, longer incubation up to 5-6 days were required to form black colonies. All genome-editing experiments were performed in triplicates. Mean and standard deviations were reported in this work.

### Screening for multiplexed genome editing

The crRNA expression cassette of MET25 and URA3A was subcloned into pYLXP’-AsCpf1-AscrRNA-CAN1 to result in pYLXP’-AsCpf1-AscrRNA-CAN1-MET25 and pYLXP’-AsCpf1-AscrRNA-CAN1-URA3A, respectively. The plasmids were transformed into *Y. lipolytica* po1g and po1fΔALK7::*URA3*, respectively. For the selection of *can1* and *met25* double mutants, yeast colonies were first spotted on CSM-Arg agar plates supplemented with L-canavanine. Positive colonies were replicated on MLA plates.

The screening of double mutants of *CAN1* and *URA3* was performed by growing the yeast transformation mixture in liquid CSM-Leu media supplemented with 50 mg/L uracil. After 4 days of outgrowth, cell culture was plated on CSM-Leu agar plate supplemented with 50 mg/L uracil and 1 mg/mL 5’-FOA. Positive colonies were spotted on CSM-Arg agar plate with L-canavanine.

Disruption of three genes was then investigated by subcloning of the crRNA-URA3A-U8 expression cassette into pYLXP’-AsCpf1-AscrRNA-CAN1-MET25 to yield pYLXP’-AsCpf1-AscrRNA-CAN1-MET25-URA3A-U8. This construct was transformed into strain po1fΔALK7::*URA3*. Transformation mixture was cultured in liquid CSM-Leu media supplemented with 50 mg/L uracil for 4 days. Cell culture was then plated on CSM-Leu agar plate with 50 mg/L uracil and 1 mg/mL 5’-FOA. After 2 days of incubation, 5’-FOA resistant colonies were screened on CSM-Arg+L-canavanine agar plate. The canavanine-resistant colonies were subsequently streaked on MLA plates to identify MET25 mutations. All genome-editing experiments were performed in triplicates. Mean and standard deviations were reported in this work.

### Y. lipolytica transformation and colony-PCR

Yeast transformation was performed using the lithium acetate method as previously described (Gaillardin, Ribet et al. 1985). Transformation mixture was diluted and plated on proper agar plates and incubated for 2-3 days until colonies appeared. Yeast colonies were picked up and boiled in 10 µL 0.02 M sodium hydroxide for 10 min at 95 °C. Primers and Q5 PCR mix (New England Biolabs, USA) were added to perform colony PCR. PCR products were Sanger sequenced to identify in-del mutations.

## Acknowledgements

Dr. Zhiliang Yang was supported by the Bill & Melinda Gates Foundation under grant no. OPP1188443. Harley Edward was supported by National Science Foundation under grant no. 1805139. Dr. Xu would like to thank the helpful discussion and suggestions with Prof. Jef Boeke and the Boeke lab members at NYU Langone Medical School.

## Author contributions

PX conceived and supervised the work. ZLY and HE performed the studies. PX and ZLY wrote the manuscript with editing from HE.

