## Supplemental Tables and Figures for "CRISPR-Cas12a/Cpf1-assisted precise, efficient and multiplexed genome-editing in *Yarrowia lipolytica*"

### Supplementary Tables

**Supplementary Table S1.** Oligos, primers and synthetic genes used in this work.

| Primer name | Primer sequence (5'-3') |
|---|---|
| AsCpf1_F | ccgaccagcactttttgcagtactaaccgcagacacagttcgagggctttaccaacc |
| AsCpf1_R | GGACAGGCCATGGAACTAGTCGGTACCTTACTTTTTCTTTTTTGCCTGGCCGGCC |
| AvrII_Fwd | GCATCCCTAAATTTGATGAAAGC |
| SalI_Rvs | GTTACATCCTTTTATCAGACATAGTC |
| TEF_F | GGGTATAAAAGACCACCGTCCCC |
| XPR2_R | CCGTTGTAGGCAACAGCGTTGGG |
| Can1crR | GATATCGGGTAGTAAAGACGGATCTACAAGAGTAGAAATTAAC |
| Can1crF | CCGTCTTTACTACCCGATATCTTTTTTTTACTAGTTCCATGGCCTG |
| Can1crR-TEF | ATCTACAAGAGTAGAAATTACTGCGGTTAGTACTGCAAAAAGTGCTGGTC |
| Can1crF-TEF | TAATTTCTACTCTTGTAGATCCGTCTTTACTACCCGATATCACTAGTTCCATGGCCTGTC |
| AvrII-SCR1-F | GGCATCCCTAAATTTGATGAAAGCCTAGGCCCCAGTTGCAAAAGTTGACACA |
| tRNA-Gly-R | CGGCGCATTAGAGGTATTTTTCAAAGTAGCCCAGTGCAGAGTCCAG |
| tRNA-Gly-F | CTGGACTCTGCACTGGGCTACTTTGAAAAATACCTCTAATGCGCCG |
| tRNA-Gly-Spe-R | GCAACGTGGGGACAGGCCATGGAACTAGTAAAAAAAAATCTACAAGAGTAGAAATTA |
| Can1-5sRNA-R | ATCAGACATAGTCGACAAAAAAAAGATATCGGGTAGTAAAGACGGATCTACAAGAGTAGA |
| Can1-chkF | CGCACAAGGAAAATTACAACACC |
| Can1-chkR | GCCACCCAGAACTCGACTTCGCC |
| Can1-seqF | CCGGTCTCTTCATTGGTACC |
| URA3A-crRR | CCTTTCTGTCCTCGAACAGGAAATCTACAAGAGTAGAAATTAGACGAGCT |
| URA3A-crRF | GATTTCCTGTTCGAGGACAGAAAGGGCCGGCATGGTCCCAGCCTCCTCGC |

| | |
|---|---|
| URA3A-crRR-U8 | CAAAAAAAACTTTCTGTCCTCGAACAGGAAATCTACAAGAGTAGAAATTAGAC GAGCT |
| URA3A-crRF-U8 | GATTTCCTGTTCGAGGACAGAAAGTTTTTTTTGGCCGGCATGGTCCCAGCCTCC TCGC |
| URA3B-crRR | CGGCCTGCGAGCTGGTGCCGAGATCTACAAGAGTAGAAATTAGACGAGCT |
| URA3B-crRF | GATCTCGGCACCAGCTCGCAGGCCGGCCGGCATGGTCCCAGCCTCCTCGC |
| URA3-seqF | GTGTGCATGATCAAGACCC |
| MET25-crRR | GACGTTGATGCCGTAGGTCTTATCTACAAGAGTAGAAATTA |
| MET25-crRF | GTAGATAAGACCTACGGCATCAACGTCGGCCGGCATGGTCCCAGCCTC |
| MET6A-crRR | CCGGCAGGGCCGTCTCGGAGAGGATCTACAAGAGTAGAAATTAGACGAGCT |
| MET6A-crRF | GATCCTCTCCGAGACGGCCCTGCCGGCCGGCATGGTCCCAGCCTCCTCGC |
| MET6A-crRR-U8 | CCAAAAAAAAGGCAGGGCCGTCTCGGAGAGGATCTACAAGAGTAGAAATTAG ACGAGCT |
| MET6A-crRF-U8 | GATCCTCTCCGAGACGGCCCTGCCTTTTTTTTGGCCGGCATGGTCCCAGCCTCC TCGC |
| MET6B-crRR | CCCAAGGATATCCGAATCAACCATCTACAAGAGTAGAAATTAGACGAGCT |
| Met6B-crRF | GATGGTTGATTCGGATATCCTTGGGGCCGGCATGGTCCCAGCCTCCTCGC |
| MET2(A)-crRF | GTAGATCTGGTCCGAGACCAGTCCCGAGGCCGGCATGGTCCCAGCCTC |
| MET2(A)-crRR | GACGTTGATGCCGTAGGTCTTATCTACAAGAGTAGAAATTA |
| MET2(B)-crRF | GTAGATCCACGCTCTTACCGGTTCCGCGGCCGGCATGGTCCCAGCCTC |
| MET2(B)-crRR | TCGGGACTGGTCTCGGACCAGATCTACAAGAGTAGAAATTA |

**Supplementary Table S2.** Plasmids and Strain genotype used in this study.

| Strains or plasmids | Description | Reference |
|---|---|---|
| **Strains** | | |
| *E. coli* NEB 5α | *fhuA2* Δ(*argF-lacZ*)*U169 phoA glnV44 Φ80 Δ(lacZ)M15 gyrA96 recA1 relA1 endA1 thi-1 hsdR17* | New England Biolabs |
| *Y. lipolytica* po1g | MATa, *leu2*-270, *ura3*-302::URA3, *xpr2*-3 | Lab stock |
| *Y. lipolytica* po1f | MATa *ura3*-302 *leu2*-270 *xpr2*-322 *axp2*-deltaNU49 *XPR2*::*SUC2* | Lab stock |
| *Y. lipolytica* po1fΔALK7 | *Y. lipolytica* po1fΔ*ALK7*::*URA3* | Lab stock |
| **Plasmids** | | |
| pYLXP' | Cloning vector | Lab stock |
| pY010 | pcDNA3.1-hAsCpf1 | Addgene[54] |
| pYLXP'-AsCpf1 | pYLXP' carrying AsCpf1 gene | This work |
| pYLXP'-AscrRNA-CAN1 | pYLXP' carrying crRNA of CAN1 under TEF-intron promoter with ribozymes | This work |
| pSCR1'-tRNA$^{Gly}$-CAN1 | pYLXP' carrying crRNA of CAN1 under SCR1'-tRNA$^{Gly}$ promoter | This work |
| pYLXP'-AscrRNA-CAN1-TEF | pYLXP' carrying crRNA of CAN1 under TEF-intron promoter without ribozymes | This work |
| pYLXP'-AscrRNA-5sRNA-CAN1 | pYLXP' carrying crRNA of CAN1 under 5s rRNA promoter | This work |
| pYLXP'-AscrRNA-URA3A | pYLXP' carrying crRNA-URA3A of URA3 under TEF-intron promoter with ribozymes | This work |
| pSCR1'-tRNA$^{Gly}$-URA3A | pYLXP' carrying crRNA-URA3A of URA3 under SCR1'-tRNA$^{Gly}$ promoter | This work |
| pYLXP'-AscrRNA-URA3B | pYLXP' carrying crRNA-URA3B of URA3 under SCR1'-tRNA$^{Gly}$ promoter | This work |
| pYLXP'-AscrRNA-URA3A-U8 | pYLXP' carrying crRNA-URA3A-U8 of URA3 under TEF-intron promoter with ribozymes | This work |
| pYLXP'-AscrRNA-MET6A | pYLXP' carrying crRNA-MET6A of URA3 under TEF-intron promoter with ribozymes | This work |

| | | |
|---|---|---|
| pYLXP'-AscrRNA-MET6B | pYLXP' carrying crRNA-MET6B of URA3 under TEF-intron promoter with ribozymes | This work |
| pYLXP'-AscrRNA-MET6A-U8 | pYLXP' carrying crRNA-MET6A-U8 of URA3 under TEF-intron promoter with ribozymes | This work |
| pYLXP'-AsCpf1-AscrRNA-CAN1 | pYLXP' carrying AsCpf1 gene and crRNA of *CAN1* | This work |
| pYLXP'-AsCpf1-SCR1'-tRNA$^{Gly}$-CAN1 | pYLXP' carrying AsCpf1 gene and crRNA of *CAN1* under SCR1'-tRNA$^{Gly}$ promoter | This work |
| pYLXP'-AsCpf1-AscrRNA-CAN1-TEF | pYLXP' carrying AsCpf1 gene and crRNA of *CAN1* under TEF-intron promoter without ribozymes | This work |
| pYLXP'-AsCpf1-AscrRNA-5sRNA-CAN1 | pYLXP' carrying AsCpf1 gene and crRNA of *CAN1* under 5s rRNA promoter | This work |
| pYLXP'-AsCpf1-AscrRNA-URA3A | pYLXP' carrying AsCpf1 gene and crRNA-URA3A | This work |
| pYLXP'-AsCpf1-SCR1'-tRNA$^{Gly}$-URA3A | pYLXP' carrying AsCpf1 gene and crRNA-URA3A under SCR1'-tRNA$^{Gly}$ promoter | This work |
| pYLXP'-AsCpf1-AscrRNA-URA3B | pYLXP' carrying AsCpf1 gene and crRNA-URA3B | This work |
| pYLXP'-AsCpf1-AscrRNA-URA3A-U8 | pYLXP' carrying AsCpf1 gene and crRNA-URA3A-U8 | This work |
| pYLXP'-AsCpf1-AscrRNA-MET25 | pYLXP' carrying AsCpf1 gene and crRNA-MET25 | This work |
| pYLXP'-AsCpf1-AscrRNA-MET6A | pYLXP' carrying AsCpf1 gene and crRNA-MET6A | This work |
| pYLXP'-AsCpf1-AscrRNA-MET6B | pYLXP' carrying AsCpf1 gene and crRNA-MET6B | This work |
| pYLXP'-AsCpf1-AscrRNA-MET6A-U8 | pYLXP' carrying AsCpf1 gene and crRNA-MET6A-U8 | This work |

**Tale S3**. Colonies count and editing efficiency for CAN1.

| CAN marker | Canavanine resistant colonies/Total colonies | | | | |
|---|---|---|---|---|---|
| **Promoter** | Experiment 1 | Experiment 2 | Experiment 3 | Average (%) | Standard deviation (%) |
| TEF-Intr-Ribo | 12/16 | 13/16 | 10/16 | 72.92 | 9.55 |
| TEF-Intr | 10/10 | 10/10 | 8/10 | 93.30 | 11.52 |
| SCR1'-tRNA-Gly | 9/10 | 9/10 | 8/10 | 86.63 | 5.74 |
| 5s rRNA | 3/10 | 3/10 | 2/10 | 26.50 | 5.63 |

**Tale S4**. Colonies count and editing efficiency for URA3.

| URA3 marker | FOA resistant/total colonies | | | | |
|---|---|---|---|---|---|
| | Experiment 1 | Experiment 2 | Experiment 3 | Average (%) | Standard deviation (%) |
| Control | 1/20 | 1/20 | 2/20 | 6.67 | 2.89 |
| TEF-Intr-Ribo | 12/12 | 8/8 | 10/10 | 100.00 | 0.00 |
| Control | 2/20 | 1/20 | 1/20 | 6.67 | 2.89 |
| SCR1'-tRNA-Gly | 14/14 | 10/10 | 9/12 | 91.67 | 14.43 |
| Control | 1/20 | 1/20 | 1/20 | 5.00 | 0.00 |
| TEF-intr-Ribo-U8 | 10/10 | 10/10 | 9/10 | 96.67 | 5.77 |
| Control | 1/20 | 1/20 | 1/20 | 5.00 | 0.00 |
| TEF-intr-Ribo | 11/11 | 10/10 | 7/10 | 90.00 | 17.32 |

**Tale S5**. Colonies count and editing efficiency for MET25/MET2/MET6.

| MET marker | Black colonies/Total colonies | | | | |
|---|---|---|---|---|---|
| | Experiment 1 | Experiment 2 | Experiment 3 | Average (%) | Standard deviation (%) |
| Met25 | 13/20 | 11/20 | 12/20 | 60.00 | 5.00 |
| Met2a | 8/10 | 7/10 | 7/10 | 73.33 | 5.77 |
| Met2b | 5/10 | 6/10 | 5/10 | 53.33 | 5.77 |
| Met6a | 16/20 | 14/20 | 15/20 | 75.00 | 5.00 |
| Met6b | 8/20 | 7/20 | 9/20 | 40.00 | 5.00 |
| Met6aU8 | 10/20 | 11/20 | 12/20 | 55.00 | 5.00 |

**Tale S6** Colonies count and editing efficiency for duplex and triplex genetic markers.

| Multiplexed targets | FOA resis or black + Can resis colonies/total colonies | | | | |
|---|---|---|---|---|---|
| | Experiment 1 | Experiment 2 | Experiment 3 | Average (%) | Standard deviation (%) |
| CAN1-MET25 | 16/20 | 18/20 | 16/20 | 83.33 | 5.77 |
| CAN1-URA3 | 14/20 | 16/20 | 15/20 | 75.00 | 5.00 |
| CAN1-URA3-MET25 | 10/20 | 8/20 | 7/20 | 41.67 | 7.64 |

### Supplementary Figures

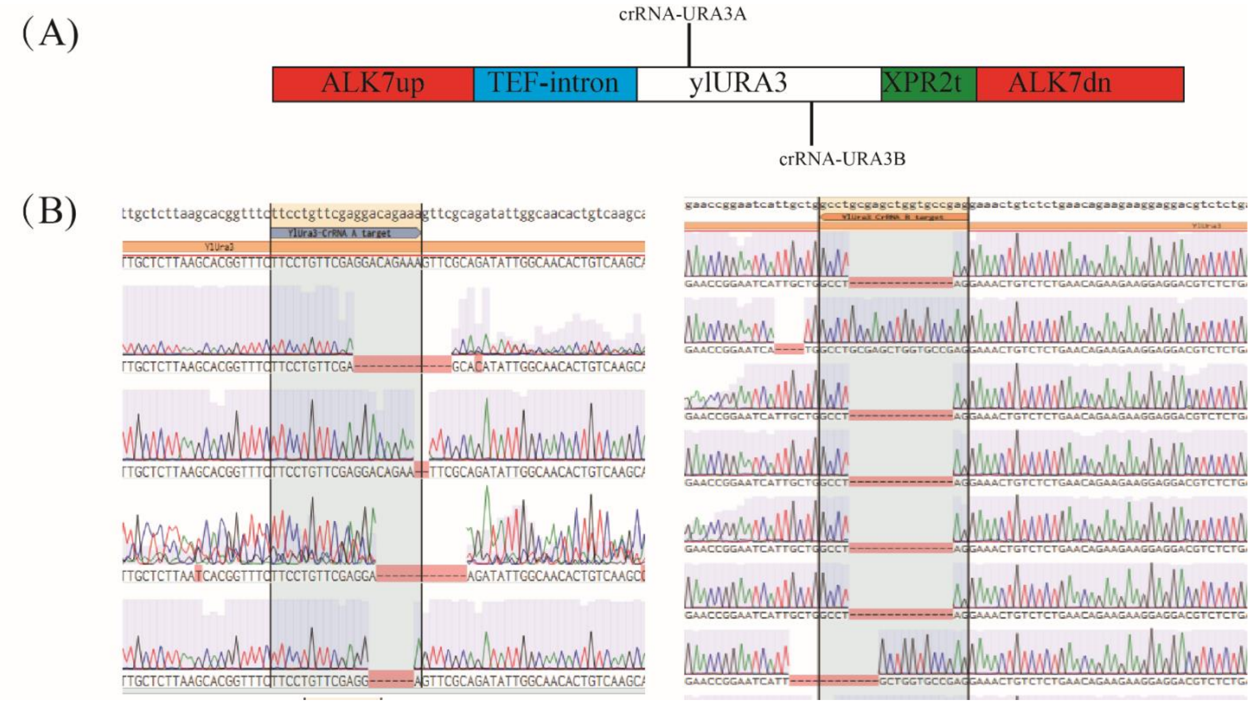

**Supplementary Figure S1**. Genomic loci for URA3 (A) and sanger DNA-sequencing for crRNA targeting either to the sense strand or the antisense strand (B).

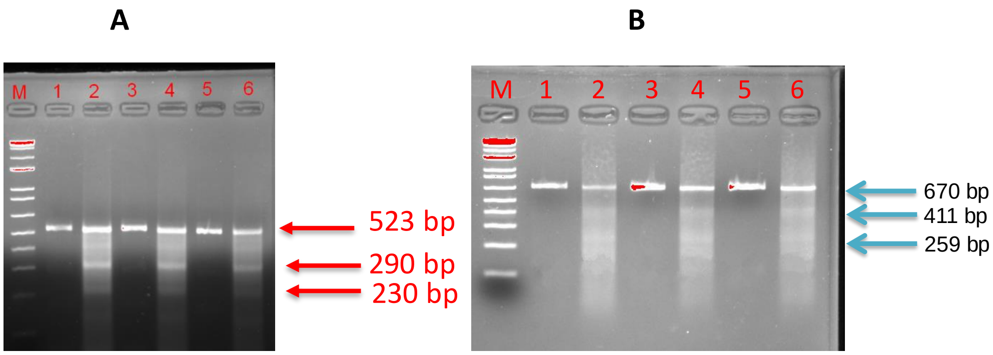

**Supplementary Figure S2.** Indel mutation detection with the endonuclease activity of Surveyor kits for MET25 (**A**) and MET6 (**B**). T7E endonuclease will cut double strand DNA when there is at least one base pair mismatch. PCR samples were first denaturalized and later the annealed heteroduplex DNAs will be digested with T7E to detect indel mutations. (**A**) For MET25, the original PCR band is about 523 bp; positive indel mutations will generate band around 290 bp and 230 bp. Lane M: 1 kb plus DNA ladder; Lane 1, 3, 5: Homodimer of MET25 mutant a, b,c; Lane 2, 4, 6: Heterodimer of MET25 mutants digested with Surveyor kits. (**B**) For MET6, the original PCR band is about 670 bp; positive indel mutations will generate band around 411 bp and 259 bp. Lane M: 1 kb plus DNA ladder; Lane 1, 3, 5: Homodimer of MET6 mutant a, b,c;; Lane 2, 4, 6: Heterodimer of MET6 mutants digested with T7E.

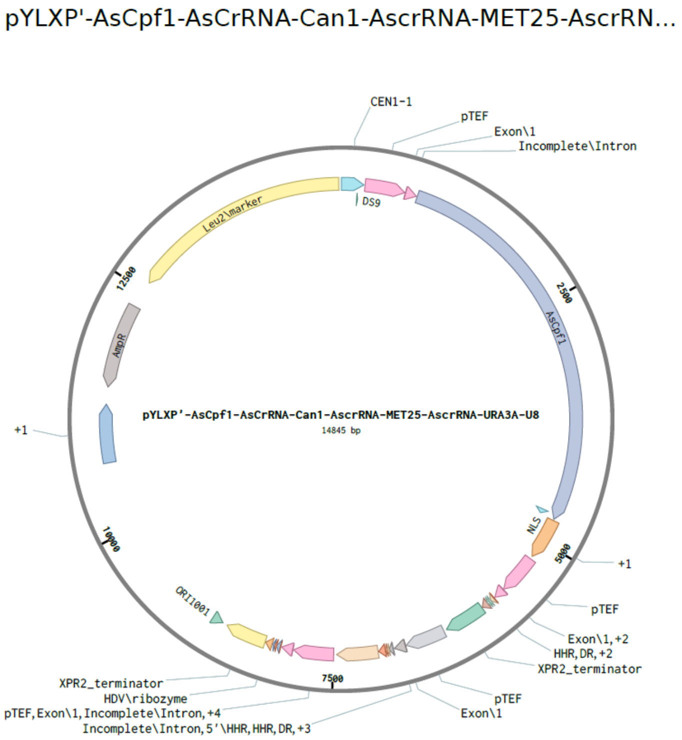

**Supplementary Figure S3.** Plasmid map and genetic configuration for three crRNA (URA3, MET25 and CAN1) arrays constructed in the lab.

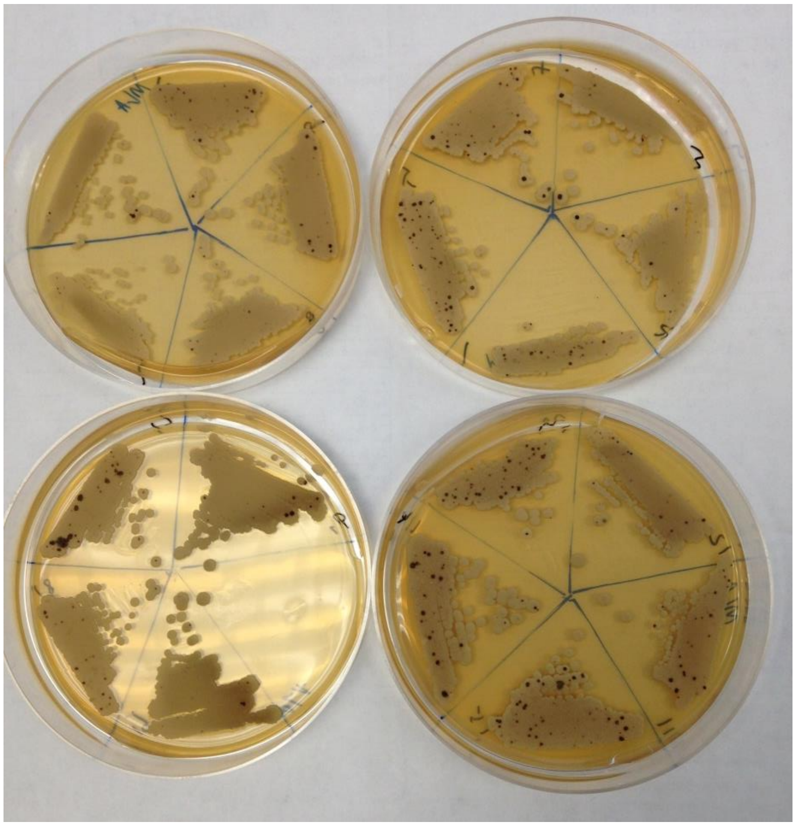

**Supplementary Figure S4**. Genetic screening for duplex and triplex genome-editing in Y. lipolytica. Cells were grown either on MLA plate, or CSM-Arg+Canavanin plate or CSM+5FOA plates. The number of black colonies indicate the incidence of successful mutation of MET25 in *Y. lipolytica*.
